# A Gene Replacement Humanization Platform for Rapid Functional Testing of Clinical Variants in Epilepsy-associated *STXBP1*

**DOI:** 10.1101/2021.08.13.453827

**Authors:** Kathryn McCormick, Trisha Brock, Matthew Wood, Lan Guo, Kolt McBride, Christine Kim, Lauren Resch, Stelian Pop, Chandler Bradford, Preston Kendrick, Jennifer A. Lawson, Adam Saunders, Sarah McKeown, Ingo Helbig, Matthew N. Bainbridge, Christopher E Hopkins

## Abstract

**Purpose:** Functional evidence is a pillar of variant interpretation according to ACMG guidelines. Functional evidence can be obtained in a variety of models and assay systems, including patient-derived tissues and iPSCs, in vitro cellular assays, and in vivo assays. Here we evaluate the reliability and practicality of variant interpretation in the small animal model, *C*. *elegans*, through a series of experiments evaluating the function of syntaxin binding protein, STXBP1, a well-known causative gene for Early infantile epileptic encephalopathy 1 (EIEE1).

**Methods:** Using CRISPR, we replaced the coding sequence for unc-18 with the coding sequence for the human ortholog *STXBP1*. Next, we used CRISPR to introduce precise point mutations in the human *STXBP1* coding sequence, reflecting three clinical categories (benign, pathogenic, and variants of uncertain significance (VUS)). We quantified 26 features of the resulting worms’ movement to train Random Forest (RF) and Support Vector Machines (SVM) machine learning classifiers on known pathogenic and benign variants. We characterized the classifiers, and then used the behavioral data from the VUS-expressing animals to predict the categorization of the VUS.

**Results:** Whereas knock-out worms without unc-18 are severely impaired in motor function, worms expressing *STXBP1* in its place have restored motor function. We produced worms with *STXBP1* variants previously classified by ACMG criteria, including 25 benign variants, 32 pathogenic, and 24 variants of uncertain significance (VUS). Using either SVM or RF classifiers, we were able to obtain a sensitivity of 0.84-0.97 on known benign and pathogenic strains. By comparing multiple ML classification methods, we were able to classify 9 of the VUS as functionally abnormal, suggesting that these VUS are likely to be pathogenic.

**Conclusions:** We demonstrate that automated analysis of a small animal system is an effective, scalable, and fast way to understand functional consequences of variants in *STXBP1*, one of the most common causes of genetic epilepsies and neurodevelopmental disorders.

## Introduction

Due to the low cost of DNA sequencing as a tool in disease diagnostics, large volumes of variant data are being generated and aggregated in databases such as ClinVar ^1,2^. Interpretation of these variants by ACMG guidelines falls into one of 5 categories: benign (B), likely benign (LB), variants of uncertain significance (VUS), likely pathogenic (LP) and pathogenic (P). VUS are problematic as they are frequently interpreted as non-diagnostic by clinicians^3^, which can impede genetic diagnosis for individuals and thereby access to appropriate medical care. Acquiring a diagnosis is critical for obtaining effective therapy, medical reimbursement, and fully informed comprehension for the family or individual affected by a genetic diagnosis. One of the strategic visions set forth by the National Human Genome Research Institute (NHGRI) is to vastly increase clinical relevance predictions, rendering the term “VUS” obsolete by 2030^4^.

Missense variants are one of the most challenging types of variants to classify accurately^5^. As of August 2020, there were 340,635 missense variants in the ClinVar database assessed with the 5 classifications of clinical significance (B, LB, VUS, LP, P), with 73% of them being VUS (Figure 1A). Prediction of variant pathogenicity can be accomplished in many ways. *In silico* tools are valuable because they are inexpensive to generate and can be used to precompute every possible variant in the genome. However, even best in class *in silico* predictions have a ROC of only ∼80%^6^. Functional characterization in an animal model is considered strong evidence for variant interpretation and further, animal models can then be used to test therapeutics.

**Fig. 1.**
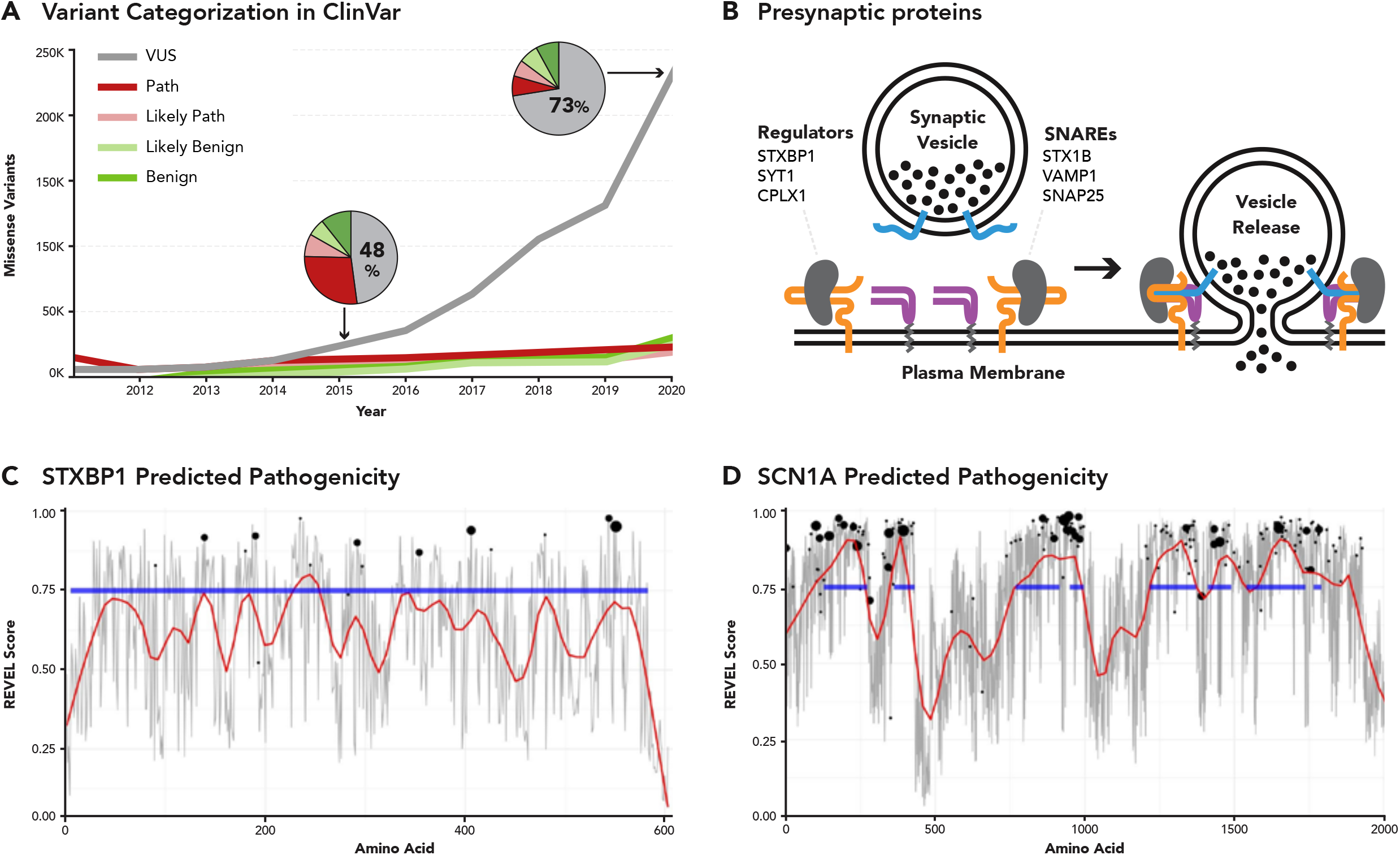
Growing need for classification in reported variants of STXBP1. Figure 1: a) Number of missense variants in ClinVar that are VUS (gray), Pathogenic (red), Likely Pathogenic (pink), Likely Benign (light green), or Benign (green) as a function of time. b) The role of STXBP1, the protein of focus in this paper, in coordinating vesicular release. Associated proteins are listed. c) In silico predicted pathogenicity from REVEL, mean score per amino acid (gray), and local regression smoothing (red) across STXBP1, with super family domain (blue), and known pathogenic variants (black dots); dot size is relative to number of pathogenic variants at that locus. d) As in c, but for SCN1A.

Mouse models are widely used as human analogs but are challenging to generate and have a high cost per data point. Like rodents, the *C*. *elegans* nematode is a multicellular organism with a variety of differentiated tissues (gut, nerves, muscle, etc.) that interact and give rise to complex behaviors. *C*. *elegans* is an extremely well studied model organism: all somatic cells and their lineages are known, behavioral phenotypes are easily quantifiable, and the application of genetic tools and methodologies is advanced, enabling the single-copy locus specific insertion of large genetic cargo, as is performed in this body of work. By humanizing *C*. *elegans*, that is replacing the worm ortholog with the human gene at the native locus, we can model human clinical variants and assess the effect on the animal (Figure 2 A, B). These unique properties of fast and affordable gene editing, automated behavioral phenotyping, cellular imaging, and molecular characterization make the *C*. *elegans* nematode a uniquely suitable organism for use in functional studies of clinical variants.

**Fig. 2.**
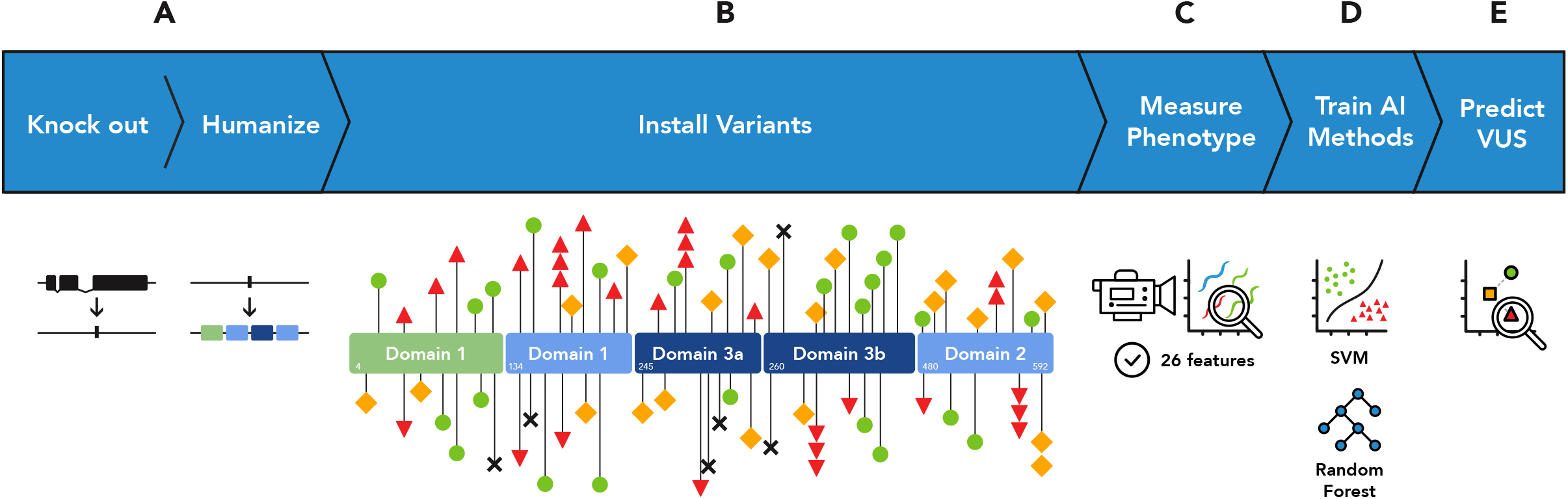
Schematic representation of the experiments performed. A) The native homolog of STXBP1, *unc-18*, was removed from the genome in a full deletion knockout. Subsequently, a codon optimized coding sequence encoding human STXBP1 is inserted into the same genomic location. B) Individual human variants were created in the STXBP1-expressing animals. The functional domains of STXBP1 as determined via crystallography are depicted, and the location of individual variants are marked with shapes. Red triangles represent pathogenic missense variants in our training data set, black X’s represent pathogenic truncating variants, green circles represent benign variants, yellow squares represent VUS. C) The generated strains were automatically assessed for 26 phenotypic features characterizing the animals’ movement and morphology. D) Two machine learning models, random forest (RF) and support vector machines (SVM) were trained on the resulting dataset. E) Models were used to sort VUS into functionally normal and abnormal groups, representing functional predictors of pathogenicity.

Mutations in *STXBP1* are one of the most common single-gene causes of developmental and epileptic encephalopathy, a neurodevelopmental disorder in which more than 80% of individuals diagnosed have seizure^7^. While initially identified in individuals with severe early-onset epilepsy^8^, the phenotypic spectrum has significantly broadened since with a large number of individuals presenting with broad neurodevelopmental features^9^. Infantile Spasms, typically starting around the age of six months, are one of the most common seizure types in *STXBP1*-related neurodevelopmental disorders. In a recent cohort from a large pediatric healthcare network^10^, *STXBP1* was the only genetic etiology significantly associated with infantile spasms and epileptic spasms. While seizures represent a major issue in infants, developmental issues with a preponderance of communication issues are the main concern in older individuals and adults. The full phenotypic range of *STXBP1*-related disorders is currently unknown, and major questions regarding natural history and features in adulthood remain unanswered.

*STXBP1* encodes a synaptic vesicle protein critically involved in vesicle release. Homozygous Stxbp1 knock-out mice demonstrate completely abolished synaptic function, emphasizing the pivotal role of STXBP1^11^. STXBP1 has a wide range of functions, including the transport of Syntaxin 1A (STX1A), a vesicular release protein, from the soma to the synapse via binding at STX1A’s Habc subdomain^12^. STXBP1 also enables Syntaxin to form SNARES at the synaptic terminal and fuse with the membrane for vesicular release via binding at a second site, the N-terminal^13–18^ (Figure 1B).

Models of STXBP1 dysfunction exist across the entire spectrum of research animals and cell lines, including yeast, flies, worms, mice, HEK293 cell culture, and differentiated fibroblasts, underlining its importance. However, two animal models, mouse and *C*. *elegans*, have recently emerged as the main organisms of research for the widespread study of variant function by several groups^16–19^. Using these organisms, researchers have recently been able to provide evidence for differing disease mechanisms, including haploinsufficiency, in which a protein truncation or nonsense variant results in a loss of translated functional protein, and dominant negative function, in which missense mutations result in misfolded proteins that aggregate with and disrupt the function of wild-type copies^7,8,16,17,20,21^. Only recently, protein stabilization strategies have been identified and are currently being assessed in clinical trials^22^.

Therapeutic discovery for STXBP1 disorders has been undertaken using structure-based *in silico* screening of 255,780 compounds paired with characterization of 17 hits in HEK293 cell culture and 3 hits in *C*. *elegans* model^17^. This approach was successful in identifying 3 commercially available compounds that were capable of rescuing STXBP1 protein levels as well as functional spontaneous and evoked neurotransmission in two disease associated variants, G544D and R406H. Intriguingly, of the three compounds explored on two variants, each variant had a unique compound that was most capable of restoring protein stability and function, indicating that variant-specific therapies may be an important approach to STXBP1 disease.

STXBP1 is distinct from many of the other common epilepsy genes in that it does not have multiple distinct functional domains, which makes variant interpretation difficult, and nearly half of *STXBP1* ClinVar variants are VUS. Ensemble *in silico* prediction tools such as REVEL^23^ and BayesDel^24^ show that for ion channel genes such as *SCN1A*, regions of high predicted pathogenicity overlay onto superfamily domain structure. However, the predicted pathogenicity for variants in *STXBP1* is relatively uniform, meaning that no regions can be regarded categorically as likely benign (Fig 1C and D). This analysis suggests that functional modeling of *STXBP1* variants will be necessary to resolve VUS.

## Materials and methods

### Creation of humanized wildtype animals

The coding sequence for the most abundantly expressed isoform of *STXBP1* (isoform a) was extracted from the UniProt database (https://www.uniprot.org/). The sequence was codon-optimized for transgene expression in *C*. *elegans*. Three synthetic introns were introduced, and the sequence was further optimized for enabling splicing specific only to the introduced introns. The sequence was introduced into the native unc-18 locus using CRISPR gene editing. Two sgRNA sites in unc-18 were selected, one in the 5’UTR of unc-18 and the other in the last exon of unc-18. A plasmid was designed to provide donor homology sequences that flank the outside edges of the two cut sites. Within the plasmid sequence and between the two donor homology sequences, the codon-optimized *STXBP1* sequence was introduced using standard molecular cloning techniques. Care was taken to design for elimination of the sgRNA sites to avoid recutting of the edited locus. The plasmid also contained a Hygromycin resistance cassette for selection^25^. The plasmid along with Cas9 and sgRNAs was injected into the hermaphrodite gonad to elicit transgene insertion^26^. Animals were harvested by Hygromycin B selection for transgene insertion and homozygous animals were identified^27^. Verification of the desired edit was confirmed by PCR and was followed by DNA sequencing. After confirmation of the desired edit, an additional round of genome editing was performed to remove the antibiotic selection sequence and restore the native 3’UTR sequence. A second round of PCR and DNA sequencing was performed to confirm that the desired sequence composition of the humanized *STXBP1* transgenic worm was obtained (the “hSTXBP1” strain). After strains were genotypically confirmed, we quantified mRNA expression as a validation that the variants introduced into the coding sequence were being transcribed.

### Creation of variant animals

For each variant, a set of sgRNA were selected to flank the locus of interest. A donor homology oligonucleotide (dhODN) was designed to have at least 35 base pairs of homology on the outside ends of the cut sites. In the interval between the cut sites, the DNA was recoded with a new sequence, containing both the desired amino acid change and silent mutations that block recutting. The dhODN was co-injected into the hermaphrodite gonad with the appropriate sgRNAs and Cas9 and dpy-10 co-CRISPR reagents. Animals were harvested for transgene insertion and homozygous animals were harvested. Verification of the desired edit was confirmed by PCR followed by DNA sequencing, and transgenic lines were also confirmed to be wild-type at the dpy-10 co-CRISPR locus. We found no significant differences in mRNA expression between variants and control sequences. We also performed an in silico assessment for any unintentionally introduced splice variants and confirmed that all flagged variant lines spliced as expected via sequencing.

### Deep Phenotyping Data Acquisition

Ten adult animals from bleach synchronized populations were transferred to clean NGM growth plates seeded with HB101 bacteria. A copper ring was melted into the agar to keep the worms from escaping the field of view. Worms were allowed to acclimate for 10 minutes on the plates after which videos were recorded for 10 minutes using the WormLab platform (MBF Bioscience, VT). For each variant three independent biological replicate populations were assayed. The WormTracker software (MBF Bioscience, version 2020.1.1) was used to track each worm’s movement throughout the 10 minute recording time. Tracks were repaired to account for collisions between worms and with the copper ring. Non-worm signals were removed. The WormTracker software was then used to analyze and export the worm movement behavior and morphology features.

### Machine Learning and Classifiers

For our training data we aggregated by genotype, by taking the mean across worms of the same genotype for each feature, so that each genotype only had one representative row.

The SVM was implemented using SciKit Learn version 0.24.1 in Python 2. Five-fold cross validation on the training set was performed to select the best performing learning algorithm and hyperparameters. A SVM with RBF kernel and gamma= .137, C =1 was selected. The features were normalized before training began.

Our RF model was produced using R^28^ and randomForest ^29^. We built the random forest using all available input features. To determine values for mtry and ntree we built random forests across a range of possible values and plotted the out-of-bag errors. Altering these parameters did not significantly impact performance of the RF. To determine performance we generated the fraction of out-of-bag pathogenic votes for each training genotype of our training genotypes (mtry=15, ntree=500). This vote fraction shows classification agreement among trees and can be used as a confidence score.

### Principal component analysis and Linear Discriminant analysis

Principal component analysis and Linear Discriminant analysis were performed according to standard methods and implemented in sklearn 0.24.1. We fit our PCA and LDA models using averages of each genotype’s 26 features and performed a transform. The first components of the PCA and LDA transformed data (PC1 and LD1, respectively) are the x and y values for each genotype in Figure 4.

### ClinVar Analysis

The ClinVar database was used to examine the clinical significance of missense variants as a function of time. ClinVar was queried with creation dates for variants on an August to July academic calendar (i.e., “2010/08/01 [Creation Date] : 2011/07/31 [Creation Date]”. The results were then restricted to missense variants. The number of variants in each Clinical Significance category were then recorded from the filter table at left, and plotted as a function of ending year.

## Results

### Insertion of human *STXBP1* transgene as gene replacement of the *C*. *elegans* unc-18 gene results in the rescue of loss of function activity

Often clinical variant modeling is performed as the insertion of an amino acid change into the native ortholog locus of an animal model. Yet in *C*. *elegans*, where 7 of every 10 disease-associated genes have an ortholog, the sequence identity is often near or below 50%. The sequence identity between *STXBP1* and its ortholog in *C*. *elegans*, *unc-18*, is 59%, and the number of conserved positions for established pathogenic and likely pathogenic variants listed in ClinVar is just over 80%. (n=48; n=58 respectively). This means that 17.2% (or nearly 1 in 5) of the established Pathogenic/Likely-Pathogenic variations cannot be modeled at the orthologous position in *C*. *elegans* (see supplemental table 1). To exacerbate the problem, 40% of the VUS are not conserved and could not be adequately modeled in the native protein. To circumvent this limitation, the entire gene locus was humanized by replacing the endogenous coding sequence in *C*. *elegans* with a human transgene sequence for the human *STXBP1* wild type gene.

The *STXBP1* coding sequence was inserted as a gene replacement of the orthologous *unc-18* locus using CRISPR-based gene editing techniques. The result was a genomic-integrated transgene expressed at single copy and regulated by endogenous regulatory sequences (see methods) for the native protein. Analysis of Electropharyngeogram (EPG) recordings and swimming in liquid indicated that the STXBP1 protein rescued abnormal phenotypic function measured in the unc-18 deletion strain, restoring activity to near wild-type levels (Supplemental Figure 1). STXBP1 protein expression was also confirmed via Western blot analysis.

### Creation and deep phenotyping of *C*. *elegans* encoding human clinical variants

We used CRISPR-based gene editing to introduce clinically relevant variants into the humanized gene locus. A set of 81 clinical single nucleotide human population variants listed in ClinVar were selected to be modeled in consultation with clinicians and patient advocacy groups, including 25 benign, 31 pathogenic, and 24 VUS. We classified the variants for input into our machine learning models according to standardized criteria. The benign variants were those that had a benign or likely benign designation in the ClinVar database, or those that had a high-frequency as reported in the GnomAD unaffected control population database (3 observations or a frequency of 1.19e-5). The pathogenic variants were those that had pathogenic and likely pathogenic annotations in the ClinVar database as of January 2021. To ensure loss-of-function phenotypes would be present, six of the pathogenic variants were selected from protein truncation variants that disrupt full-length expression at various positions along the primary amino acid sequence (R122X, R367X, R388X, Y140X, E302X, K308X). The variants that met the criteria for pathogenic and benign were used as a training set. The VUS were selected from ClinVar and in consultation with clinical groups at Rady Children’s Hospital and Children’s Hospital of Philadelphia (Figure 2C). The resultant strains were phenotyped using a semiautomated system in which locomotion and morphology of individual animals were video recorded and quantified via software algorithm, resulting in 26 phenotypic features (Figure 2D).

### Data Exploration and Feature Engineering

Primary data analysis and exploration indicated that no single feature could be used as a reliable indicator of functionally abnormal protein activity to delineate pathogenic and benign variants . We conducted principal component analysis and determined that the first two PCs only accounted for 64% of the variance in the data, indicating that there was very little redundancy in the data. This analysis also suggested that significant information would be lost in dimensional reduction. Thus, we performed K-means clustering using all 26 features as inputs. Plotting the K-means clustering into the dimensional space of PC1 and PC2 revealed three distinctive clusters (Supplemental Figure 2). One cluster contained 16 pathogenic, including 6 truncating mutations, as well as the unc-18 KO worm. Another cluster was mixed, containing 21 benign and 14 pathogenic variants, and the unmutated, humanized worm. A third cluster contained two pathogenic variants and the N2, wild-type worm.

### Training of Machine Learning Algorithms

In order to generate a phenotype-based decision boundary for VUS, we undertook training of two machine learning classifiers with the established pathogenic and benign variants. We used two orthogonal classifier algorithms: random forest (RF), which generates a classification status based on the majority call vote of a series of decision trees, and support vector machines (SVM), which uses training data to generate a hyperplane decision boundary that can be used to classify new inputs. The two classifiers were in good agreement, differing on their classification on only 4 of the 57 variants in the training set (Figure 3 A, B). We evaluated the performance of the classifiers by examining the confusion matrices, receiver operator characteristic plots and precision recall plots. Both classifiers performed well. The RF had an AUROC of 0.84 and an AUPRC of 0.87 whereas the SVM had an AUROC of 0.94 and an AUPRC of 0.96 (Figure 3 D, E). Both classifiers misclassified the pathogenic variants R551C, R551S, R551L, E283K, G544V, R292H, R292L, S42Y as benign. They also both misclassified the benign variants D207N, V104I, and T588I as pathogenic variants. The truncation variants were consistently classified correctly, and were among the most pathogenic according to the distance from the decision boundary (SVM) or percentage of decision tree votes (RF).

**Figure 3:**
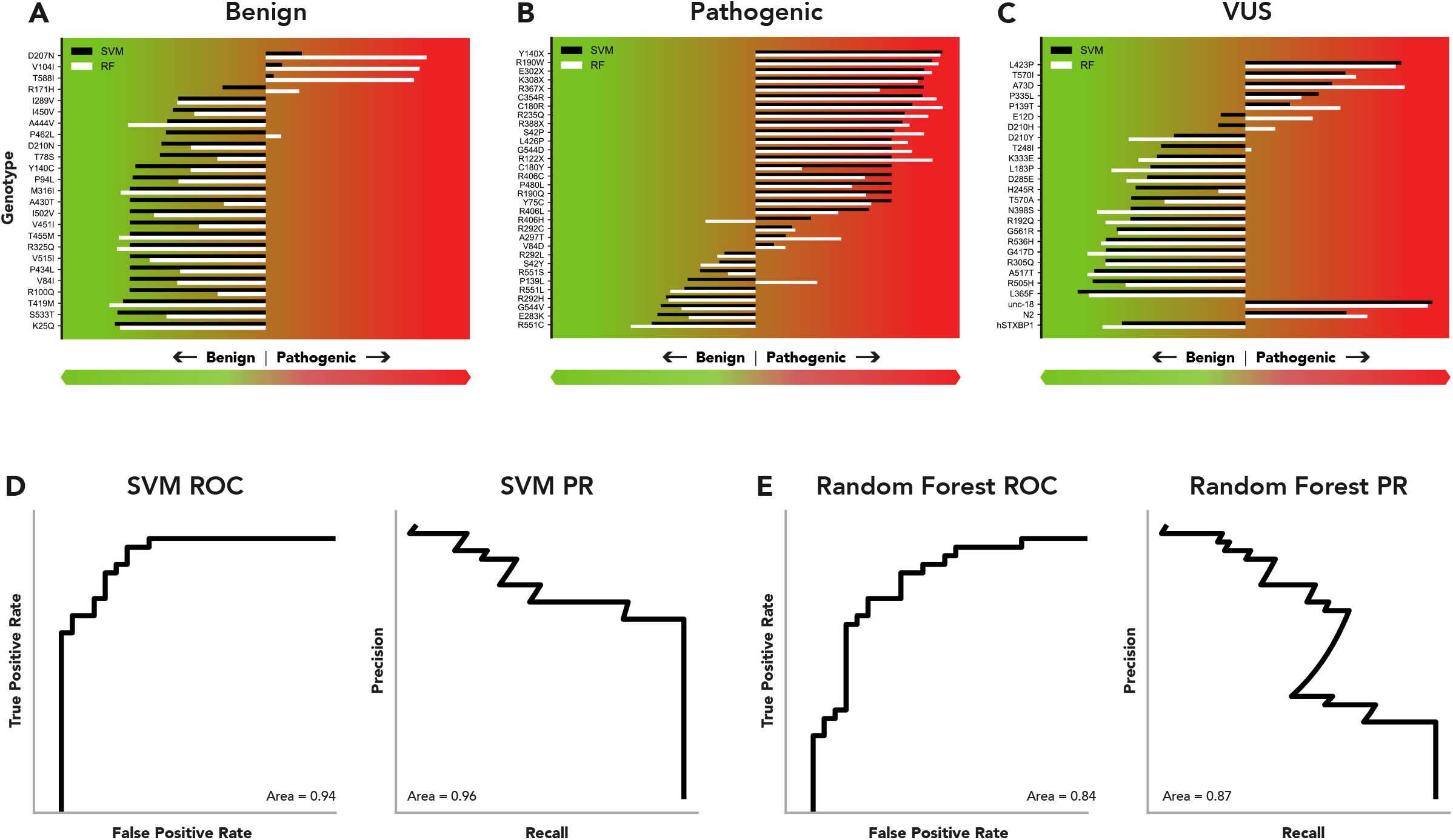
Model evaluation and functional predictions. Top row: Normalized functional scores from the SVM (black) and RF (white) model. Benign (A) , Pathogenic (B) and VUS (C) variants in our data set were evaluated by the trained models. The x axis represents the scaled distance from the decision boundary (middle point) between benign (left) and pathogenic (right) predictions, indicating confidence in the prediction. Bottom Row: Receiver Operator Characteristic (ROC) and Precision- Recall (PR) curves for SVM (D) and RF (E) methods.

### Evaluation of VUS

Of the 24 VUS evaluated by the two models, the SVM and RF both classified 6 variants as pathogenic: L423P, D597N, T570I, A73D, P335L, and P139T (Figure 3C). The classifiers disagreed on three variants (D210H, E12D, and T481I) with the RF classifier judging them as pathogenic, whereas the SVM classifier judged them as benign. All other VUS were classified as benign by both methodologies.

Further analysis of the spatial localization of the variants in the multivariate space defined by the first principal component and the first linear discriminant revealed that pathogenic variants fell along two distinct axes (Figure 4). One axis is composed of protein truncation variants and presumed loss of function variants. The unc-18 knockout worm also falls along this axis. The second axis is sparsely populated, and comprises the pathogenic variants Y75C, R406C, and the wildtype N2 worm. The VUS P335L, classified as pathogenic, also falls along this axis in the functional assay space.

**Fig 4.**
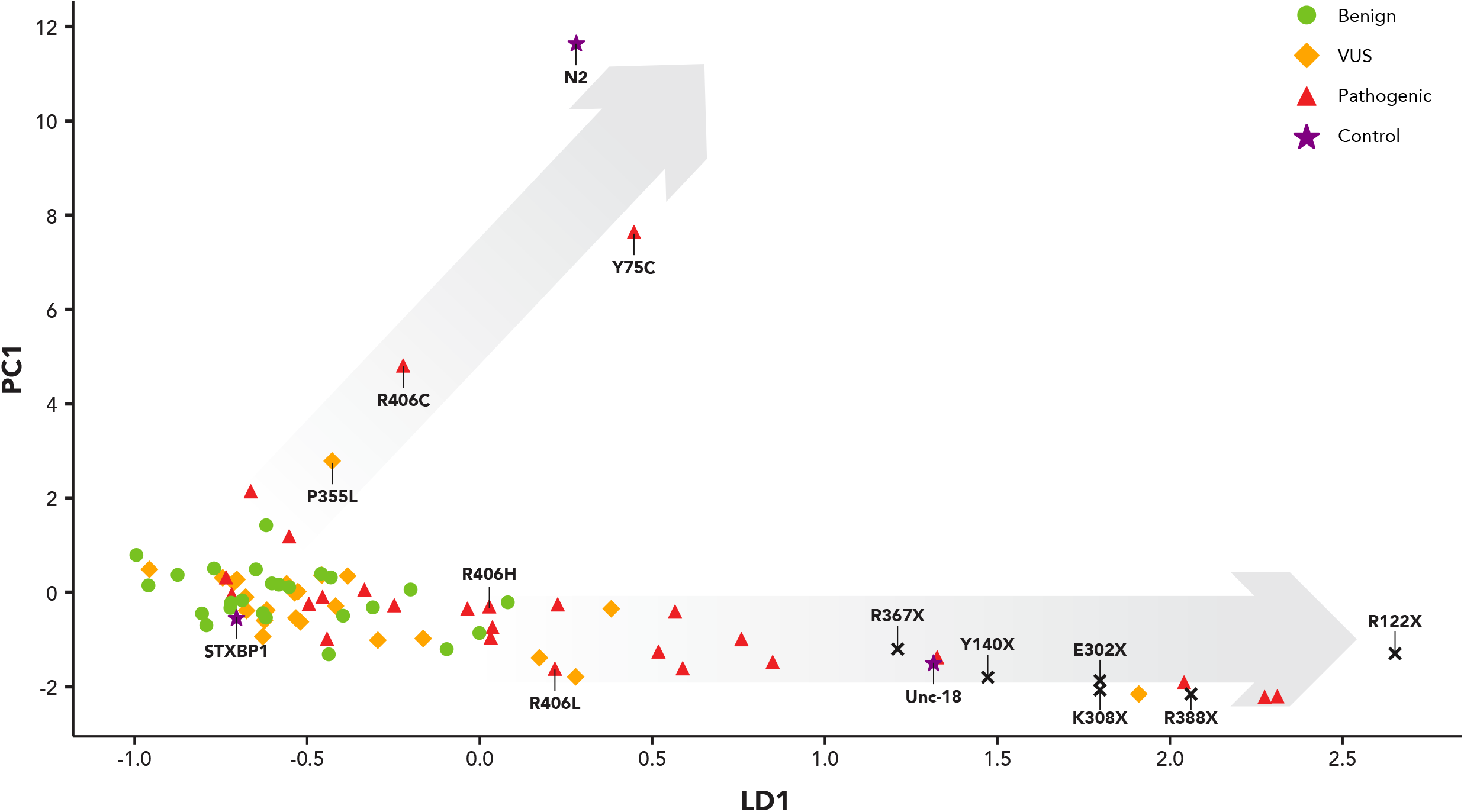
Visualization of all strains in multiparameter space. Each strain is represented by its value on the first linear discriminant (LD1) vs its value on the first principal component. Three control strains, provided for comparison, are represented by purple stars: the genetically humanized STXBP1, the *unc-18* knockout, and the wildtype N2 strain. Benign variants used in the training set are represented by green circles. Pathogenic variants used in the training set are represented by red triangles. VUS are represented by orange diamonds. Grey translucent arrows represent two discernible axes of variation in the data in which pathogenic variants are distinguishable from the genetically humanized STXBP1 strain.

## Discussion

This work demonstrates that gene-humanized animals expressing *STXBP1* can be used as a line of functional evidence in determining variant pathogenicity. The variants created and functionally assessed in this paper account for a significant proportion of variants seen in individuals with *STXBP1* related disorders, including most recurrent variants in the *STXBP1* gene. We estimate that in this work we have screened approximately 25% of all disease-causing variants in STXBP1 reported in the literature since the discovery of the *STXBP1* gene as cause of human epilepsy, reflecting the scalability of our assay system.

Early-Infantile Epileptic Encephalopathy 4 (EIEE4) is an autosomal dominant disorder which manifests in individuals heterozygous for disease causing variants within *STXBP1*^30^. The animal models herein are phenotyped in their homozygous state. The biallelic effect of the clinical variants is modeled, and thus mechanisms such as haploinsufficiency and dominant negative function cannot be resolved. In our assay, the both classifiers struggled with the mutational hotspots R551 and R292, misclassifying 3/3 and 2/3 variants respectively, however, the classifiers correctly identified all mutations at the hotspot R406. Further investigation into the mechanism of pathogenicity for variants involving these positions is needed. It is possible that our biallelic modeling results in an inability to detect dominant negative functions. Another potential limitation of our approach is the use of only one isoform. STXBP1 has two isoforms with an alternative last exon choice. The shorter isoform is 10x more common in the GTEX v7 data set and we chose to model all variants in this isoform. Thus we were unable to evaluate clinical variants in the longer, more rarely expressed isoform.

Despite the limitations, the findings presented herein represent the development of a fast and efficient platform for functional testing of clinically-observed variants in a disease-associated gene that can be applied in a whole animal context. Approaches in live animals may present an important alternative to deep mutational scanning in cells. Although extremely powerful and efficient, deep mutational scanning requires that the gene being analyzed results in an lethal phenotype, which may be either inherent or in an engineered genetic context. Additionally, since genes have different functions in different tissues, mutational analysis in a single cell type may not be representative of overall organismal results. *In vivo* mutational modeling, as performed here, has the additional benefit of enabling therapeutic discovery on behavioral phenotypes and allowing simultaneous discovery of mechanisms (e.g. GOF and LOF) that can be tied to molecular readouts (e.g., mRNA and protein expression). Performing the equivalent level of clinical variant modeling with germline integration of single nucleotide polymorphisms in the mouse model would require a much longer timeline before any phenotypic evaluation could be performed, and although disease modeling and mechanism research is advancing in the zebrafish, the field is currently rate-limited by the extreme difficulty of generating point mutations via homology directed repair (HDR). Performing CRISPR based HDR editing in C. elegans is routine and can be performed on timelines relatively close to cell culture. To our knowledge, such an extensive variant analysis in a disease-associated gene has not been performed in any other model organism. The results of this study could inform future study designs in, e.g., iPSCs, which would provide an important validation.

This VUS analysis platform could be applied to other genes where functional assessment is required, but is not amenable to deep mutational approaches. We suggest that candidate genes for the approach taken herein have several qualities: (1) homology between the human and C. elegans genes is >50%; (2) The human disease gene has 2 or fewer high-homology orthologs in C. elegans; and (3) the overall size of the human coding sequence isis <5KB. A preliminary analysis of the allelic variants with genotype-phenotype relationships cataloged in the On-line Mendelian Inheritance in Man (OMIM) database indicated that 2058 of 4689 (44.5%) would meet these criteria. As a result, a large number of human genes are likely to be amenable to the same gene humanization technique applied to STXBP1 and variant pathogenicity can be tested in a whole animal format.

Discrimination between loss-of-function (LOF) and gain-of-function (GOF) variants may be key in appropriately choosing therapeutics for individuals with disease-causing variants. Finding both GOF and LOF variants is common in dosage-sensitive genes including many genetic etiologies implicated in the epilepsies, such as *SCN2A* or *SCN8A* ^31,32^. In KCNQ2-associated diseases, the use of antiepileptic drugs effective for LOF variants of *KCNQ2* is associated with poor outcomes in patients with a GOF variant^33^. Differentiating between GOF and LOF variants can give key insights into the prognosis expected for an individual.

Interestingly, though most pathogenic and truncating variants occupied a similar phenotypic feature space as the native *unc-18* knockout in our assay, a few pathogenic variants and pathogenic-assigned VUS occupied a space more similar to the wildtype worms in our data, which may indicate that they have increased synaptic release compared to the unmutated humanized worm. Among these was the P335L variant, which was included in our data set as a VUS due to having a conflicting designation in ClinVar. However, many lines of evidence point to this mutation having a hypersecretion phenotype with increased neurotransmitter release ^17,18,20,34–36^. This suggests an intriguing possibility that this *C. elegans in vivo* assay may be capable of discriminating LOF and GOF mutations. More work and additional lines of evidence are necessary to confirm this possibility.

Despite the accuracy of our assay, the humanized worm did not acheieve a complete rescue of the unc-18 KO phenotype (Figure 4, Supplementary Figure 2) and a higher level of discrimination between the benign and pathogenic categories might be possible if complete rescue of function was achieved. STXBP1 is a presynaptic protein whose primary role is to provide coordinated release of neurotransmitters at the synapse. STXBP1 directly binds syntaxin and influences syntaxin’s binding interaction with the two other key proteins necessary for promoting vesicle fusion, synaptobrevin and SNAP25. The lack of complete rescue with the human transgene may be due to imperfect fit between STXBP1 and the *C*. *elegans* subunits of the SNARE complex. Since the common ancestor of *C*. *elegans* and humans was 500 million years ago, the presynaptic proteins have coevolved as their sequence drifted over time. Future work to introduce multiple human proteins into the *C*. *elegans* synapse may increase the predictive power of this animal model system.

## Conclusions

We describe a fully humanized *C*. *elegans* model for rapid modelling of the pathogenic effects of variants in *STXBP1*, a common cause of genetic epilepsy. We demonstrate that our genetic models can be used within automated phenotypic analysis and machine learning workflows to distinguish disease-associated pathogenic variants from benign variants, thereby providing a scalable assay to understand variant pathogenicity and VUS resolution. We used the disease modeling platform to functionally characterize 24 VUS and present evidence that 6 of the VUS are functionally abnormal.

## Supplementary material

Supplementary material is available online.

## Data availability

xxx

## Acknowledgments

xxx

## Author Information

xxx

## Ethics Declaration

xxx

