## Supplemental figures, legends, and methods for "A Gene Replacement Humanization Platform for Rapid Functional Testing of Clinical Variants in Epilepsy-associated *STXBP1*"

[1] InVivo Biosystems. Eugene, OR, 97402, USA.

[2] Codified Genomics, LLC, Houston, TX, 77004, USA.

[3] Division of Neurology, Children's Hospital of Philadelphia, Philadelphia, PA, 19104, USA.

[4] The Epilepsy NeuroGenetics Initiative (ENGIN), Children's Hospital of Philadelphia, Philadelphia, PA, 19104, USA.

[5] Department of Biomedical and Health Informatics (DBHi), Children's Hospital of Philadelphia, Philadelphia, PA, 19146, USA.

[6] University of Pennsylvania, Neuroscience Program, Philadelphia, PA, 19104, USA.

[7]Rady Children's Institute for Genomic Medicine, San Diego, CA, 92123, USA.

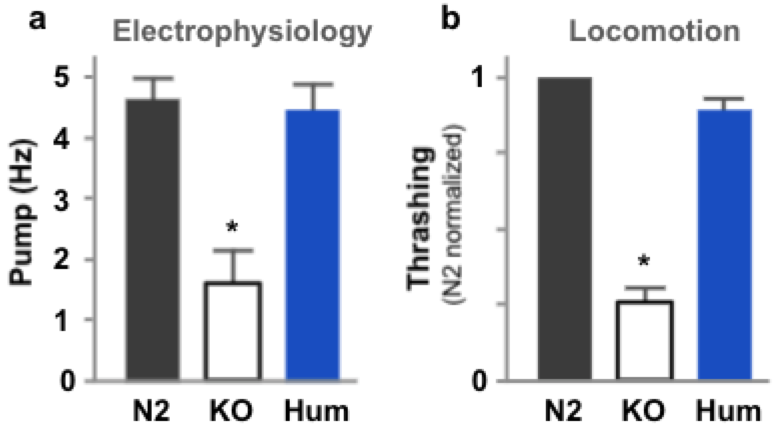

**Supplemental Figure 1**: Phenotypic charaterization of wildtype (N2), *unc-18* knockout (KO), and STXBP1 gene replacement humanization (Hum) worms using two assays. (a) Electrophysiological characterization on pharyngeal pumping rate stimulated with serotonin. (b) Locomotion rate in liquid media (thrashing), where the locomotory cycle is scored in cycles/sec (Hz), and normalized to the wildtype N2 worm.

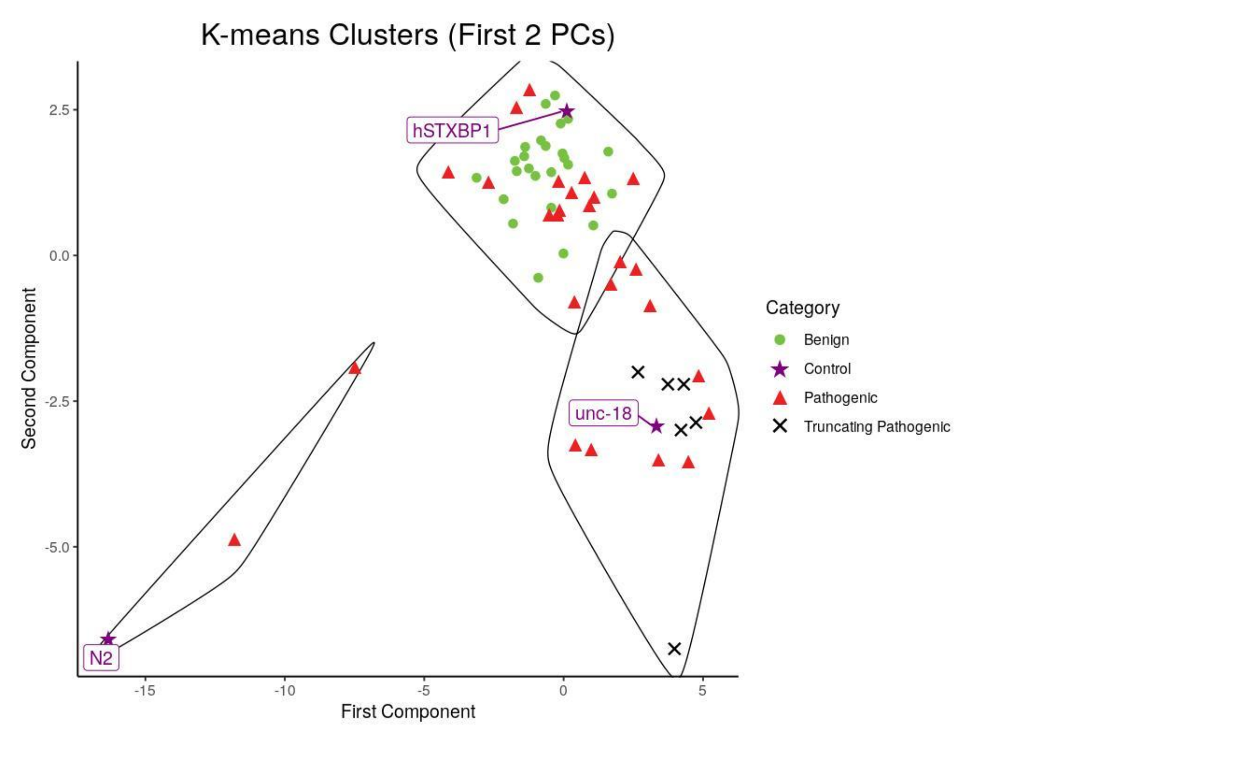
K-means Clusters (First 2 PCs)
Second Component
First Component
2.5
0.0
-2.5
-5.0
-15
-10
-5
0
5
hSTXBP1
unc-18
N2
Category
Benign
Control
Pathogenic
Truncating Pathogenic

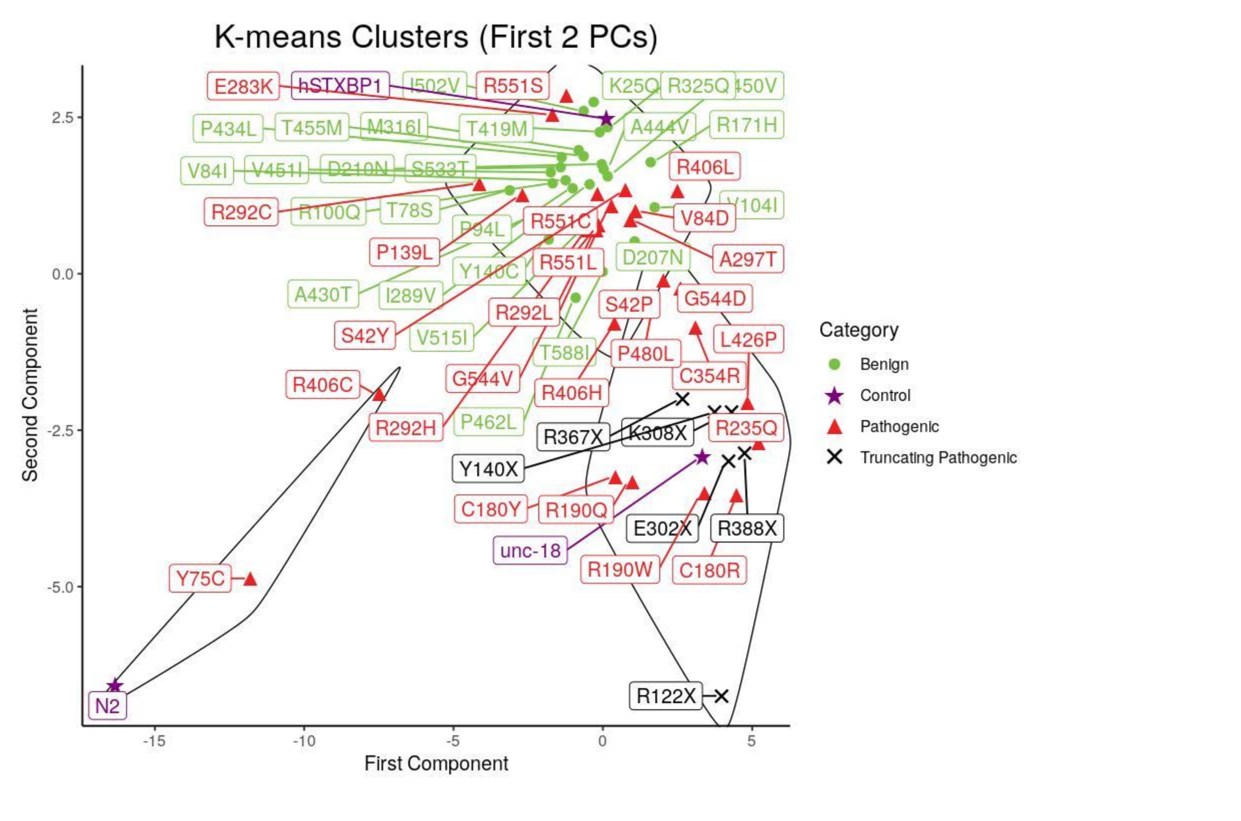
K-means Clusters (First 2 PCs)
Second Component
First Component
2.5
0.0
-2.5
-5.0
-15
-10
-5
0
5
E283K
hSTXBP1
R551S
R325Q
P434L
T455M
M316I
T419M
A444V
R171H
V84I
V451I
D210N
S533T
R406L
R292C
R100Q
T78S
V84D
P94L
R551C
P139L
R551L
D207N
A297T
Y140C
A430T
I289V
S42P
G544D
R292L
S42Y
V515I
L426P
T588I
P480L
G544V
C354R
R406C
R406H
R235Q
P462L
R292H
R367X
K308X
Y140X
C180Y
R190Q
E302X
R388X
unc-18
R190W
C180R
Y75C
R122X
N2
Category
Benign
Control
Pathogenic
Truncating Pathogenic

**Supplemental Figure 2.** K-means clustering of Benign and Pathogenic variants, visualized in the space defined by PC1 and PC2. (A) Without individual labels (b) With labels for each variant used in classification.

Supplemental Methods.

K-means clustering

The data for known pathogenic/benign genotypes was scaled and centered using Caret[1]. K-means was then performed with R[2] and the kmeans() function. To determine the number of clusters two metrics were used:

First the standard elbow method was used by plotting the total within-cluster sum of squares for 2 to 12 clusters and looking for the “elbow” or the point where the sum of squares levels off and we start to get diminishing returns [RFE: https://www.mdpi.com/2571-8800/2/2/16]. This was determined to be between 3 and 5 clusters. Next we determined cluster purity by examining the “label” (e.g. Pathogenic or Benign) of variants when known. Cluster uniformity was calculated as the mean difference in the ratios between pathogenic and benign across all clusters.

abs(pathogenic_count / total - benign_count / total)

We plotted the mean score across 2 to 12 clusters for this metric and looked for a similar inflection point as with the elbow method. This confirmed that three clusters was the correct number for this data.

To visualize k-means clusters we reduced the data down to two dimensions using PCA (R core team, princomp() with cor=True). The first two principal components for each

genotype were plotted using ggplot2[3] using the cluster assignments from the k-means analysis. To improve the labels and other aesthetics we used ggforce[4], ggrepel[5] and ggalt[6].

VUS variants were visualized by “layering” them on top of the k-means clusters generated with known benign/pathogenic variants without retraining k-means. To accomplish this the distance from each VUS to each cluster centroid was calculated using the same method used during k-means clustering. Taking the nearest centroid for each VUS gave a cluster association with the previously generated clusters.

**Column Definitions for Supplemental Table 1.**

**Amino Acid Genotype**: Change in amino acid at the protein level

**hg19 coordinates**: Variation at the genomic level, registered with hg19

**category**: The category using in our training or classification data. The benign variants were those that had a benign or likely benign designation in the ClinVar database, or those that had a high-frequency as reported in the GnomAD unaffected control population database (3 observations or a frequency of 1.19e-5). The pathogenic variants were those that had pathogenic and likely pathogenic annotations in the ClinVar database as of January 2021. All others were VUS.

**C. elegans Conservation**: The Amino acid location of the conserved locus in the C. elegans unc-18 gene.  Loci without conserved amino acids are marked “Not Conserved”.

**Scaled distance from SVM boundary**: measures the distance of each genotype from the hyperplane decision boundary.

**oob_rf_path_vote**: Fraction of out-of-bag trees that predict pathogenic (training data only)

**prediction_rf_path_vote**: Fraction of trees that predict pathogenic (VUS/controls only)

**K-means Cluster:** The k-means cluster assigned to this genotype. Data was aggregated by genotype and features were scaled to be in similar ranges before clustering was run. The total number of clusters (3) was chosen in advance. (Pathogenic, Benign, Control only)

**nearest_centroid**: Given the three centroids calculated as part of k-means, this value is the closest centroid measured by calculating the minimum sum of squares from the available centroids for the given genotype. (VUS only)

1. Kuhn, M. caret: Classification and Regression Training. *Astrophys. Source Code Libr.* ascl:1505.003 (2015).
2. R Core Team. *R: A language and environment for statistical computing*. (R Foundation for Statistical Computing, 2018).
3. Wickham, H. *ggplot2: Elegant Graphics for Data Analysis*. (Springer International Publishing, 2016). doi:10.1007/978-3-319-24277-4.
4. Pedersen, T. L. *ggforce: Accelerating 'ggplot2'.* (2020).
5. Slowikowski. *ggrepel: Automatically Position Non-Overlapping Text Labels with 'ggplot2'*. (2020).
6. Bob Rudis, Bolker, B. & Schulz, J. *ggalt: Extra Coordinate Systems, 'Geoms', Statistical Transformations, Scales and Fonts for 'ggplot2'.* (2017).
