## Supplemental Table 1 for "A Gene Replacement Humanization Platform for Rapid Functional Testing of Clinical Variants in Epilepsy-associated *STXBP1*"

| Amino Acid Genotype | hg19 coordinates | **category** | C. elegans conservation | scaled distance from SVM boundary | SVM Predicted Category | oob_rf_path_vote | prediction_rf_path_vote | RF Predicted Category | K means cluster | nearest_centroid |
|---|---|---|---|---|---|---|---|---|---|---|
| K25Q | 9:130413917A>C | benign | Not Conserved | -1.109425665 | benign | 0.1104972376 | | benign | 1 | |
| S533T | 9:130444735G>C | benign | 615 | -1.090485394 | benign | 0.2346368715 | | benign | 1 | |
| T419M | 9:130438929C>T | benign | 500 | -1.049266775 | benign | 0.08196721311 | | benign | 1 | |
| R100Q | 9:130422361G>A | benign | Not Conserved | -1.000838272 | benign | 0.3714285714 | | benign | 1 | |
| V84I | 9:130422312G>A | benign | Not Conserved | -1.000443779 | benign | 0.2631578947 | | benign | 1 | |
| P434L | 9:130438974C>T | benign | Not Conserved | -1.000198499 | benign | 0.2711864407 | | benign | 1 | |
| V515I | 9:130442517G>A | benign | Not Conserved | -1.000137395 | benign | 0.1896551724 | | benign | 1 | |
| R325Q | 9:130434340G>A | benign | Not Conserved | -1.000049268 | benign | 0.1021505376 | | benign | 1 | |
| T455M | 9:130440714C>T | benign | Not Conserved | -0.999950754 | benign | 0.1073446328 | | benign | 1 | |
| V451I | 9:130439024G>A | benign | 532 | -0.9999442952 | benign | 0.3210526316 | | benign | 1 | |
| I502V | 9:130442478A>G | benign | Not Conserved | -0.9998515478 | benign | 0.2011173184 | | benign | 1 | |
| A430T | 9:130438961G>A | benign | 511 | -0.999800495 | benign | 0.3879781421 | | benign | 1 | |
| M316I | 9:130432222G>A | benign | Not Conserved | -0.999666532 | benign | 0.1123595506 | | benign | 1 | |
| P94L | 9:130422343C>T | benign | Not Conserved | -0.979008424 | benign | 0.2666666667 | | benign | 1 | |
| Y140C | 9:130423474A>G | benign | 220 | -0.9585847037 | benign | 0.2994011976 | | benign | 1 | |
| T78S | 9:130420717C>G | benign | Not Conserved | -0.7780992527 | benign | 0.3705882353 | | benign | 1 | |
| D210N | 9:130427575G>A | benign | 290 | -0.766683012 | benign | 0.3 | | benign | 1 | |
| P462L | 9:130440735C>T | benign | 543 | -0.7331436882 | benign | 0.5411764706 | | pathogenic | 1 | |
| A444V | 9:130439004C>T | benign | 525 | -0.7231237469 | benign | 0.1318681319 | | benign | 1 | |
| I450V | 9:130439021A>G | benign | 531 | -0.6829606455 | benign | 0.3089005236 | | benign | 1 | |
| I289V | 9:130430429A>G | benign | Not Conserved | -0.6441331564 | benign | 0.2634408602 | | benign | 1 | |
| R171H | 9:130425566G>A | benign | 251 | -0.31804917 | benign | 0.5891891892 | | pathogenic | 1 | |
| T588I | 9:130453114C>T | benign | Not Conserved | 0.05852334023 | pathogenic | 0.8961748634 | | pathogenic | 1 | |
| V104I | 9:130422372G>A | benign | 184 | 0.1198205456 | pathogenic | 0.9111111111 | | pathogenic | 1 | |
| D207N | 9:130427566G>A | benign | Not Conserved | 0.265044275 | pathogenic | 0.9297297297 | | pathogenic | 1 | |
| R551C | 9:130444788C>T | pathogenic | 633 | -0.7637804794 | benign | 0.1666666667 | | benign | 1 | |
| E283K | 9:130430411G>A | pathogenic | 363 | -0.7211745801 | benign | 0.3216374269 | | benign | 1 | |
| G544V | 9:130444768G>T | pathogenic | 626 | -0.696535068 | benign | 0.3020833333 | | benign | 1 | |
| R292H | 9:130430439G>A | pathogenic | 372 | -0.6580377087 | benign | 0.2670454545 | | benign | 1 | |
| R551L | 9:130444789G>T | pathogenic | 633 | -0.522982557 | benign | 0.2717391304 | | benign | 1 | |
| P139L | 9:130423471C>T | pathogenic | 219 | -0.4984621321 | benign | 0.664893617 | | pathogenic | 1 | |
| R551S | 9:130444788C>A | pathogenic | 633 | -0.4083361016 | benign | 0.4262295082 | | benign | 1 | |
| S42Y | 9:130416031C>T | pathogenic | 124 | -0.2669015167 | benign | 0.3529411765 | | benign | 1 | |
| R292L | 9:130430439G>T | pathogenic | 372 | -0.2289990692 | benign | 0.3976608187 | | benign | 1 | |
| V84D | 9:130422313T>A | pathogenic | Not Conserved | 0.1351571317 | pathogenic | 0.5795454545 | | pathogenic | 1 | |
| A297T | 9:130430453G>A | pathogenic | 377 | 0.2220403159 | pathogenic | 0.7287234043 | | pathogenic | 1 | |
| R292C | 9:130430438C>T | pathogenic | 372 | 0.2692022653 | pathogenic | 0.6063829787 | | pathogenic | 1 | |
| R406H | 9:130438189G>A | pathogenic | 487 | 0.4077143683 | pathogenic | 0.3659793814 | | benign | 1 | |
| R406L | 9:130438189G>T | pathogenic | 487 | 0.8324409002 | pathogenic | 0.7208121827 | | pathogenic | 1 | |
| Y75C | 9:130420708A>G | pathogenic | 157 | 0.9994238917 | pathogenic | 0.8090452261 | | pathogenic | 2 | |
| R190Q | 9:130425623G>A | pathogenic | 270 | 0.9998383048 | pathogenic | 0.7955801105 | | pathogenic | 3 | |
| P480L | 9:130440789C>T | pathogenic | 561 | 0.9999609855 | pathogenic | 0.7575757576 | | pathogenic | 3 | |
| R406C | 9:130438188C>T | pathogenic | 487 | 0.9999609869 | pathogenic | 0.7923497268 | | pathogenic | 2 | |

| Amino Acid Genotype | hg19 coordinates | **category** | C. elegans conservation | scaled distance from SVM boundary | SVM Predicted Category | oob_rf_path_vote | prediction_rf_path_vote | RF Predicted Category | K means cluster | nearest_centroid |
|---|---|---|---|---|---|---|---|---|---|---|
| C180Y | 9:130425593G>A | pathogenic | 260 | 0.9999751813 | pathogenic | 0.623655914 | | pathogenic | 3 | |
| R122X | 9:130423419C>T | pathogenic | Not Conserved | 1.000029138 | pathogenic | 0.9731182796 | | pathogenic | 3 | |
| G544D | 9:130444768G>A | pathogenic | 626 | 1.000044772 | pathogenic | 0.9179487179 | | pathogenic | 3 | |
| L426P | 9:130438950T>C | pathogenic | 507 | 1.000365669 | pathogenic | 0.9076086957 | | pathogenic | 3 | |
| S42P | 9:130416030T>C | pathogenic | 124 | 1.020377425 | pathogenic | 0.9502762431 | | pathogenic | 3 | |
| R388X | 9:130438134C>T | pathogenic | Not Conserved | 1.079665508 | pathogenic | 0.9114583333 | | pathogenic | 3 | |
| R235Q | 9:130428485G>A | pathogenic | 315 | 1.09606993 | pathogenic | 0.9615384615 | | pathogenic | 3 | |
| C180R | 9:130425592T>C | pathogenic | 260 | 1.152978534 | pathogenic | 1 | | pathogenic | 3 | |
| C354R | 9:130435490T>C | pathogenic | 435 | 1.227059953 | pathogenic | 0.9836065574 | | pathogenic | 3 | |
| R367X | 9:130435529C>T | pathogenic | Not Conserved | 1.236203746 | pathogenic | 0.8323699422 | | pathogenic | 3 | |
| K308X | 9:130432196A>T | pathogenic | 388 | 1.23688119 | pathogenic | 0.9333333333 | | pathogenic | 3 | |
| E302X | 9:130432178G>T | pathogenic | 382 | 1.239764999 | pathogenic | 0.9714285714 | | pathogenic | 3 | |
| R190W | 9:130425622C>T | pathogenic | 270 | 1.297094933 | pathogenic | 0.9891891892 | | pathogenic | 3 | |
| Y140X | 9:130423475T>A | pathogenic | 220 | 1.372841477 | pathogenic | 0.9940828402 | | pathogenic | 3 | |
| L365F | 9:130435523C>T | vus | 446 | -1.232437812 | benign | | 0.082 | benign | | 1 |
| R505H | 9:130442488G>A | vus | 586 | -1.121648646 | benign | | 0.18 | benign | | 1 |
| A517T | 9:130444686G>A | vus | 598 | -1.110714632 | benign | | 0.078 | benign | | 1 |
| R305Q | 9:130432188G>A | vus | Not Conserved | -1.027923162 | benign | | 0.126 | benign | | 1 |
| G417D | 9:130438923G>A | vus | 498 | -1.024868674 | benign | | 0.078 | benign | | 1 |
| R536H | 9:130444744G>A | vus | 618 | -1.0245287 | benign | | 0.114 | benign | | 1 |
| G561R | 9:130444818G>A | vus | Not Conserved | -0.9453211247 | benign | | 0.16 | benign | | 1 |
| R192Q | 9:130425629G>A | vus | 272 | -0.8455663838 | benign | | 0.126 | benign | | 1 |
| N398S | 9:130438165A>G | vus | Not Conserved | -0.8432147605 | benign | | 0.104 | benign | | 1 |
| T570A | 9:130446652A>G | vus | Not Conserved | -0.8393947212 | benign | | 0.284 | benign | | 1 |
| H245R | 9:130428515A>G | vus | 325 | -0.8063605065 | benign | | 0.428 | benign | | 1 |
| D285E | 9:130430419C>A | vus | 365 | -0.7223371066 | benign | | 0.182 | benign | | 1 |
| L183P | 9:130425602T>C | vus | 263 | -0.6984025177 | benign | | 0.142 | benign | | 1 |
| K333E | 9:130434363A>G | vus | Not Conserved | -0.6493613407 | benign | | 0.214 | benign | | 1 |
| T248I | 9:130428524C>T | vus | 328 | -0.6196063176 | benign | | 0.516 | pathogenic | | 1 |
| D210Y | 9:130427575G>T | vus | 290 | -0.5247156044 | benign | | 0.188 | benign | | 1 |
| D210H | 9:130427575G>C | vus | 290 | -0.2002928376 | benign | | 0.58 | pathogenic | | 1 |
| E12D | 9:130374718G>C | vus | Not Conserved | -0.1806925037 | benign | | 0.68 | pathogenic | | 1 |
| P139T | 9:130423470C>A | vus | 219 | 0.3271569907 | pathogenic | | 0.754 | pathogenic | | 1 |
| P335L | 9:130434370C>T | vus | 416 | 0.5383948077 | pathogenic | | 0.65 | pathogenic | | 1 |
| A73D | 9:130420702C>A | vus | 155 | 0.637514805 | pathogenic | | 0.926 | pathogenic | | 1 |
| D597N | 9:130446733G>A | vus | Not Conserved | 0.6874418026 | pathogenic | | 0.654 | pathogenic | | 3 |
| T570I | 9:130446653C>T | vus | Not Conserved | 0.7357389903 | pathogenic | | 0.796 | pathogenic | | 1 |
| L423P | 9:130438941T>C | vus | 504 | 1.145875302 | pathogenic | | 0.902 | pathogenic | | 3 |
| hSTXBP1 | | Z | | -0.9070737154 | N/A | | 0.118 | N/A | 1 | |
| N2 | | Z | | 0.7435397862 | N/A | | 0.826 | N/A | 2 | |
| unc-18 | | Z | | 1.374981881 | N/A | | 0.988 | N/A | 3 | |
